# A Bioelectric Router for Adaptive Isochronous Neurostimulation

**DOI:** 10.1101/2025.01.28.635122

**Authors:** Eashan Sahai, Jordan Hickman, Daniel J Denman

## Abstract

**Objective:** Multipolar intracranial electrical brain stimulation (iEBS) is a method that has potential to improve clinical applications of mono- and bipolar iEBS. Current tools for researching multipolar iEBS are proprietary, can have high entry costs, lack flexibility in managing different stimulation parameters and electrodes, and can include clinical features unnecessary for the requisite exploratory research. This is a factor limiting the progress in understanding and applying multipolar iEBS effectively. To address these challenges, we developed the Bioelectric Router for Adaptive Isochronous Neuro stimulation (BRAINS) board.

**Approach:** The BRAINS board is a cost-effective and customizable device designed to facilitate multipolar stimulation experiments across a 16-channel electrode array using common research electrode setups. The BRAINS board interfaces with a microcontroller, allowing users to configure each channel for cathodal or anodal input, establish a grounded connection, or maintain a floating state. The design prioritizes ease of integration by leveraging standard tools like a microcontroller and an analog signal isolators while providing options to customize setups according to experimental conditions. It also ensures output isolation, reduces noise, and supports remote configuration changes for rapid switching of electrode states. To test the efficacy of the board, we performed bench-top validation of monopolar, bipolar, and multipolar stimulation regimes. The same regimes were tested *in vivo* in mouse primary visual cortex and measured using Neuropixel recordings.

**Main Results:** The BRAINS board demonstrates no meaningful differences in Root Mean Square Error (RMSE) noise or signal-to-noise ratio compared to the baseline performance of the isolated stimulator alone. The board supports configuration changes at a rate of up to 600 Hz without introducing residual noise, enabling high-frequency switching necessary for temporally multiplexed multipolar stimulation.

**Significance:** The BRAINS board represents a significant advancement in exploratory brain stimulation research by providing a user-friendly, customizable, open source, and cost-effective tool capable of conducting sophisticated, reproducible, and finely controlled stimulation experiments. With a capacity for effectively real-time information processing and efficient parameter exploration the BRAINS board can enhance both exploratory research on iEBS and enable improved clinical use of multipolar and closed-loop iEBS.

## Introduction

Electrical stimulation is one of a small number of clinically-tractable approaches to neuromodulation[1], along with focused ultrasound methods[2, 3, 4]. Electrical neuromodulation comes in a diversity of forms (transcranial direct current[5, 6], transcranial alternating current[7], trancranial magnetic[8], and intracranial[9]) and targets (peripheral nerves cochlea, spinal cord, and intracranial targets including the subthalamic nucleus, substantia nigra, motor cortex, sensory cortex[10]). Within the forms of electrical neuromodulation, intracranial electrical brain stimulation (iEBS) is a cornerstone therapeutic approach in clinical neurology and neurosurgery, particularly for treating Parkinson’s disease, with expanding indications for obsessive compulsive disorder[11, 12], dystonia[13], and epilepsy[14]. In addition to these expanding indications, clinical neural implants capable of iEBS are being tested as prospective sensory prosthetics[15, 16, 17] and for providing feedback to improve the quality and efficacy of motor brain-machine interfaces[18]. Furthermore, electrical microstimulation paradigms using low-amplitude and brief iEBS have been used for decades in animal research as a means activating small numbers of neurons. Despite widespread clinical use and its role in neuroscience research, our fundamental understanding of how iEBS affects neural circuits remains surprisingly limited. While we know that iEBS can activate varied neural elements - including somas, dendrites, and axons - the precise composition of recruited neural ensembles remains unclear[19, 20]. The general idea is that a small number of neurons in a uniform and symmetrical area around the stimulation site are activated; there is also ample evidence that the net effect of iEBS is suppressive even if some neurons in the field are briefly activated. This knowledge gap is particularly evident in clinical settings, where stimulation parameters must often be modified empirically by clinicians during patient follow-up visits[21], both to increase efficacy and to reduce side effects that arise through modulation of neural targets not intended to be modulated by the iEBS protocol. Multipolar brain stimulation, which extend iEBS to multiple spatially and temporally patterned electrical stimuli, has been proposed as a potential solution for increased iEBS control. Early efforts with small numbers of sites have shown promise[22] in providing more precise[23, 24] and efficient therapeutic outcomes[25, 26].

Current FDA-approved devices typically offer unipolar, bipolar, or limited multipolar stimulation options up to 8 contacts[27]. In practice, clinicians typically exhausting unipolar configurations before exploring more complex stimulation patterns[19]. Research devices with more contacts have been developed [28]; whether more advanced iEBS approaches enabled by such density, such as multipolar psuedo-contacts[29] and current steering[30], which have been pioneered in peripheral neuromodulation, can add specificity[31] to intracranial DBS-style devices is less clear. Investigation of the effects of advanced iEBS approaches, including complex multipolar stimulation, patterned stimulation, and other current steering paradigms faces significant technical and practical barriers in most research settings. The tools for exploring multipolar stimulation, such as clinical neurostimulators or high-end research systems can be either prohibitively expensive, inflexible in their configuration, or excessive in their complexity for basic research applications[25]. The research community currently lacks a flexible, open-source, cost-effective tool for exploring multipolar stimulation paradigms. These limitations have created a significant barrier to entry for researchers interested in exploring novel stimulation paradigms and understanding the basic principles of neural activation patterns.

To address these challenges, we developed a modular and readily reprogrammable interface – the Bioelectric Router for Adaptive Isochronous Neuro stimulation (BRAINS) board for control of electrical brain stimulation. This system enables rapid switching and multipolar stimulation while maintaining compatibility with existing monopolar and bipolar stimulation experimental frameworks. Our approach prioritizes accessibility, flexibility, and signal isolation while allowing integration with standard experimental setups.

## Methods

### Device Design and Prototyping

We designed, fabricated, and evaluated a system for adaptive isochronous neurostimulation that enables software controlled selection channel state when using multisite electrical stimulation devices. Functions of this device (figure 1A) include software selection of anodal and cathodal channels without needing to change physical connections, enabling multiple connections to anode and cathode channels, control of the state of non-used channels, and rapid switching between configurations. The BRAINS board was conceived and tested for use with silicon multi-channel electrodes (figure 1A, right, Neuronexus Technologies), but in principle can be coupled to any passive stimulation device.

**Figure 1:**
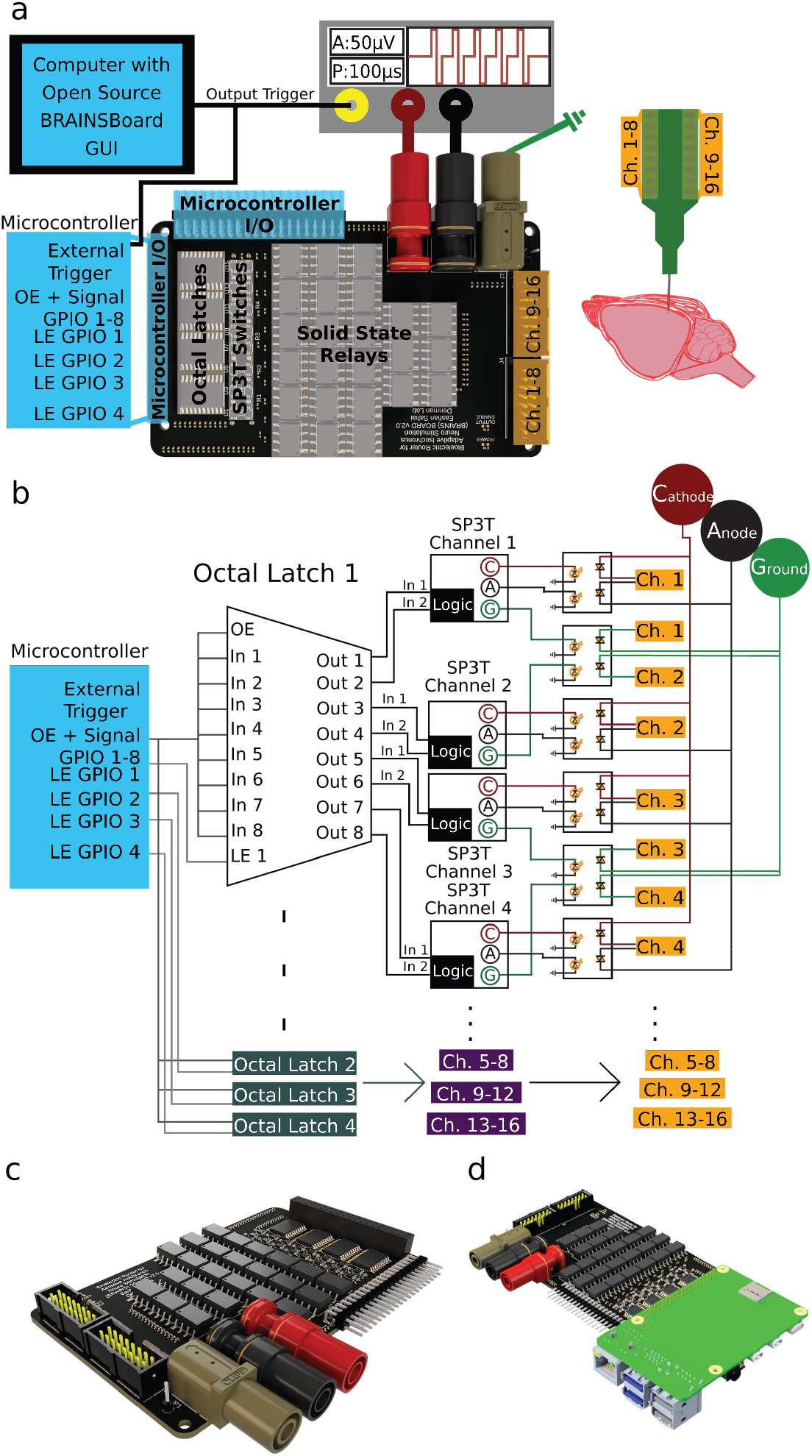
Schematic of the BRAINS board. **A**. General connection guide for full in-vivo application of the board with general part guides and microcontroller requirements with the presence of an external output trigger from a stimulating device, the key components of the board that perform all the logic (computer output to microcontroller through octal latch multiplexer throw single pull 3 throw switches) and isolation (solid state relay to split control signal from stimulator cathode/anode/ground signal connections to 16 output channels to connect to an implantable 16-channel electrode array), as well as a general setup of a in vivo invasive experiment **B**. Modeled schematic of each electronic connection within the BRAINS board demonstrating control through a microcontroller to send signals to 4 Octal latch Multiplexers to 16 Single Pull 3 throw switches and where isolation occurs through optical isolation to activate and deactivate different channels with cathode, anode, or signal ground **C**. 3D rendered BRAINS board model with connectors **D**. 3D rendered BRAINS board Model, including an integrated Raspberry Pi.

The BRAINS board provides electronic switching capabilities for 16 independent electrode channels between four states (cathode, anode, signal ground, and floating) without requiring manual intervention during experiments while maintaining signal integrity. The board’s architecture centers around four key interfaces (figure 1A): *1)* A 2×20 female header compatible with both the Raspberry Pi 4B as a direct shield and adaptable to use with any Arduino. Here, we present experiments and tests with the BRAINS board using an Arduino Pro Micro. *2)* A complementary 2×20 90° male header providing access to auxiliary pins for external sensors, actuators, grounding, power, or triggers and access to the microcontroller’s built in +5V, +3.3V, and grounds. *3)* Three standard banana connectors to connect to any analog stimulus isolator: a red connector for positive terminal, black connector for negative terminal, and a green connector for connection to a signal or building ground. There is also a pad for a 1×1 header pin that is connected to the same ground. *4)* Two standard 2×8 box connectors from Samtec (TSS-108-01-G-D) with dedicated building/signal ground connections to minimize noise, supporting flexible electrode configurations.

Signal routing and control are managed through multiple components, beginning with an octal transparent D-Type latch (Texas Instruments, CY74FCT373TSOC). This component provides stable channel selection through its latching capabilities while enabling efficient pin multiplexing. Operating at +5V, each latch interfaces directly with the microcontroller’s logic levels with a theoretical maximum 8 nanosecond delay when latching per latch **(Table 1)**. The multiplexing functionality reduces the number of required microcontroller connections to allow for scalability for future iterations and customization options beyond current intended use. The latch’s output and enable controls ensure precise timing of state changes across channels.

**Table 1.** Control Electronics Properties. Timing and electrical characteristics of the Arduino, octal latch, and SP3T switch components indicating the theoretical minimum and maximum delays for rapidly switching channels due to every piece of hardware other than the solid state relay.

| Hardware | Arduino Microcontroller |  |  | Octal Latch (74FCT373) |  |  | SP3T Switch (TS5A3357) |  |  | Units |
| --- | --- | --- | --- | --- | --- | --- | --- | --- | --- | --- |
| Characteristics | Min | Typ | Max | Min | Typ | Max | Min | Typ | Max |  |
| Delay | *** | *** | *** | 1.5 | 5.2 | 8 | 2 | 6.5 | 7 | ns |
| Input Voltage | 0 | 5 | 5.5 | 2 | - | 0.8 | 2 | - | 0.8 | V |
| Output Voltage | 0 | 5 | 5.5 | 2.4 | 3.3 | 0.55 | 2.4 | 3.3 | 0.3 | V |

To ensure deterministic state selection, an analog SP3T triple-throw switch for each potential output (Texas Instruments, TS5A3357DCUR) routes signals between the four possible states (cathode, anode, signal ground, or floating). This switch has a theoretical maximum 7 nanosecond delay to support rapid state transitions for, in the case of the current design, 16 channels independently. **(Table 1)**.

The final key component of the board design is the use of a solid state relay (IXYS Systems, PAA193STR). These relays provide optical isolation between the control circuitry and stimulation pathways to prevent introduction of noise and maintain an isolated stimulation. The optical coupling mechanism prevents unwanted electrical interference while maintaining high-voltage handling capabilities at up to 600V **(Table 2)** necessary for accommodating variable tissue and electrode impedance during current stimulation. The default configuration is ”floating”, or open; this configuration minimized the potential for unintended stimulation through an electrode not selected as anode or cathode. The board includes a circuit for status indicators: a red LED power indicator and a green LED output enable indicator that illuminates when active channel state modifications are in progress.

**Table 2.** Solid State Relay Properties: Isolation, input/output, and delay characteristics of the Solid State Relays on the BRAINS board as it theoretically affects our output signal through capacitance, leak, and limitations of the maximum Voltage that can be sent from a analog isolated stimulating device.

| Hardware | Solid State Relay (PAA193) |  |  | Units |
| --- | --- | --- | --- | --- |
| Characteristics | Min | Typ | Max |  |
| On Delay | - | - | 5 | ms |
| Off Delay | - | - | 5 | ms |
| Leak Current | - | 0.027 | 10 | μA |
| Output Capacitance | - | 50 | - | pF |
| Input Voltage | 0.9 | 1.2 | 1.5 | V |
| Output Voltage | - | - | 600 | V<sub>rms</sub> |

We designed the BRAINS board using the Electronics Design function of Autodesk Fusion360, using design files and 2-dimensional part footprints from each part’s documentation. We first designed a schematic before we placed all parts in a printed circuit board layout, ensuring through-hole vias and layers for ground and VCC along with top and bottom layers. We performed routing automatically through the built-in auto router with design rules set up for a 4-layer board with no blind or buried vias. Finally, we checked that the board passed the electrical rule check and had no airwires. We exported and packaged the gerber files, pick and place (PnP) information, and the bill of materials (BOM); original Autodesk Fusion360 CAD files with the schematic and PCB layout are written as .sch and .brd files (see the Supplement for all design and production files). We produced this board through the local third party PCB manufacturing and assembly company (Colorado PCB Assembly). The bare board production ensured that the BRAINS board was RoHS compliant. The manufacturer acquired all turnkey components listed in the BOM, and assembled the PCB as specified in the design and PnP files.

### Software design and development

We designed two primary control interfaces for the BRAINS board: an Arduino-based serial communication protocol and direct Raspberry Pi GPIO control. We first describe the Arduino-based serial protocol.

The Arduino implementation utilizes a serial communication protocol operating at a baud rate of 115200 bits per second (bps), enabling precise temporal control of electrode states through a structured command syntax. Each electrode channel can be configured to one of the four states that the SP3T switches allow: floating, cathode, anode, or signal ground. State changes are managed through the octal latch system to ensure stable transitions.

The control architecture employs multiplexing to efficiently manage the 16 channels through a minimal pin configuration. The output enable pin (Arduino Pin 16) coordinates with four latch enable pins to select specific channel groups *(1-4, 5-8, 9-12, and 13-16)*. Eight Data Input pins, connected across all octal latches, control the state transitions through a two-bit encoding scheme. This encoding determines the SP3T switch states, where the combinations of Input 1 and Input 2 (LOW-LOW: floating, HIGH-LOW: cathode, LOW-HIGH: anode, HIGH-HIGH: signal ground) define the channel configurations (figure 2A).

**Figure 2:**
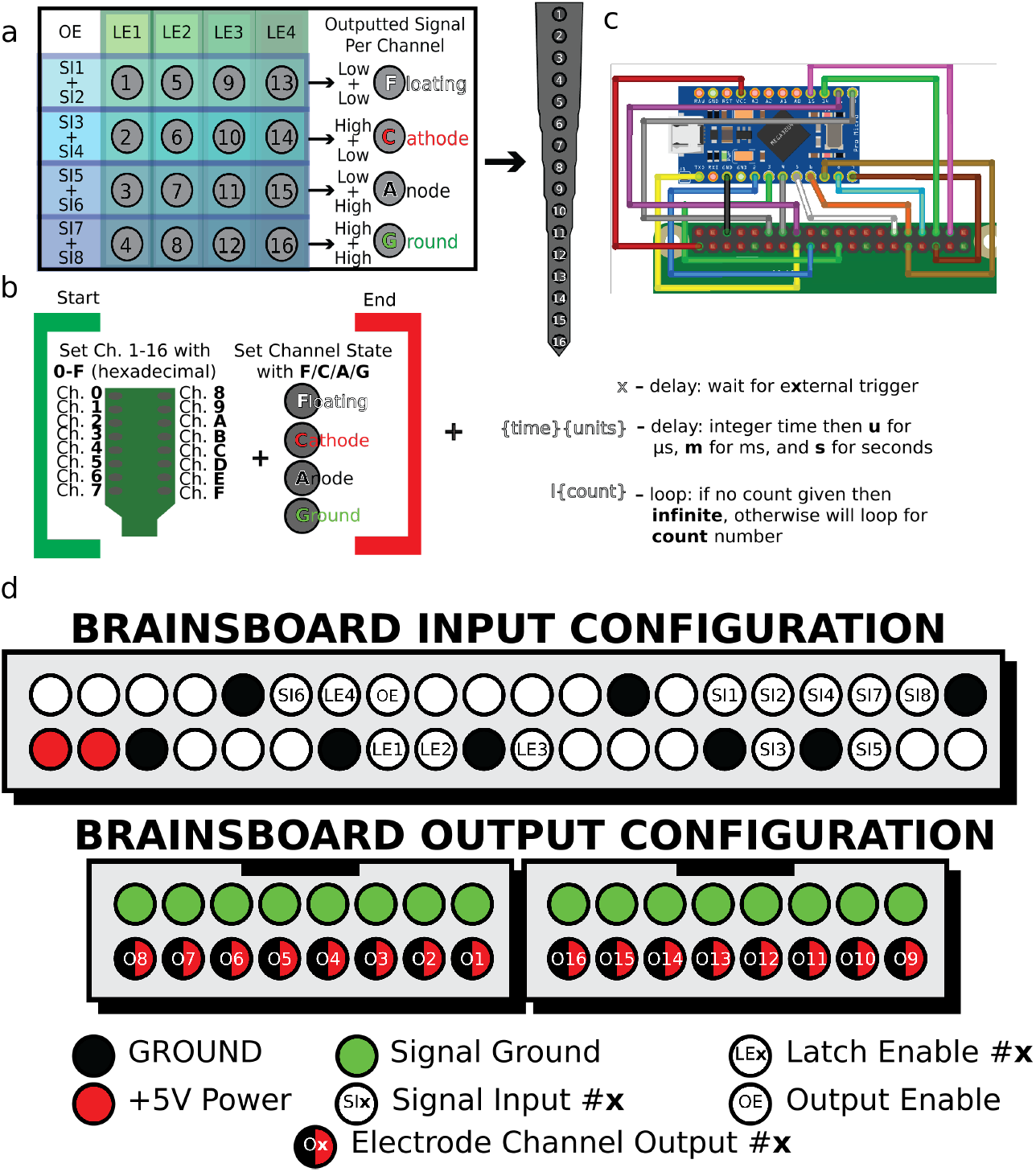
Software Procedure for the BRAINS board. **A**. Illustrated guide for input power for control devices to program individual channels with indications of what signal and latch enable outputs are required to enable a certain channel on an electrode. Each latch controls a set of 4 channels and the 8 signal pins are used in groups of 2 to determine the which of four possible channel states is selected for each channel. Any channel is modified by setting the output enable HIGH, which allows each latch to modify its 8 output states, and configuring each latch enable signal output pins to reset the HIGH/LOW state for each signal input. LOW+LOW = floating, HIGH+LOW = cathode, LOW+HIGH = anode, and HIGH+HIGH= signal ground. **B**. Basic programming guide using serial commands with indications of individual channel settings, varying forms of delays unique to electrical stimulation experiments, and how to setup loops within the framework of the Arduino Pro Micro code, with [] used to open and close setting the states, a hexadecimal value utilized to write to each individual channel, a concurrent letter corresponding to the channel state following, and key triggers between sets of [] to setup delays and loops for rapid switching purposes. 115200bps baud rate through the Arduino serial command or Arduino serial command interface (e.g., PySerial). Full code available (see Data Availability Statement). **C**. Arduino Pro Micro wiring connection schematic for setup direct to the BRAINS board **D**. Pinout for the control input 2×20 header (top) and signal output 2×8 headers (bottom) with bare minimum number of pins that are required to setup states for all individual channels as well as the exact order for channel number setup from BRAINS board to output pins. More detailed pinouts can be seen in Supplemental figure 1.

The serial command protocol implements a bracketed syntax *e.g*., ([…]) for channel configuration, supporting hexadecimal channel addressing *e.g*., *(0-9, A-F)* and state characters *e.g*., *(F, C, A, G)*. Additional control features include external trigger synchronization (via pin A0), programmable delays (in seconds, milliseconds, or microseconds), and loop functionality for repeated patterns. This protocol structure enables complex stimulation sequences while maintaining precise timing control (figure 2B). The Arduino can easily be connected using simple male to male jumper wires (figure 2C) or can be connected to any alternative microcontroller using the drawn pinout (figure 4D)

As an alternative to this serial command protocol, a Raspberry Pi interface can provide direct GPIO control through a dedicated pin mapping, eliminating serial communication overhead. This implementation is particularly advantageous for applications that benefit from wireless control, or require integration with complex input processing. The Raspberry Pi 4B’s GPIO pins 22-27 manage the BRAINS board output enable and latch enable functions, while pins 5, 6, 12, 13, 16, 17, 19, and 26 handle digital input control. This direct interface supports Python-based programming for flexible sequence generation and timing control through a precise sleep function.

Each control interface offers distinct advantages: the Arduino implementation is convenient for scenarios requiring minimal latency and integration with Windows-based instrumentation (e.g., Multi Channel Systems’ STG5 Isolated Analog Stimulator). The Raspberry Pi configuration is optimal for wireless control applications or complex input processing requirements. The choice between interfaces depends heavily on experimental requirements regarding timing precision, communication flexibility, and system integration needs. The signal ground configuration connects any channel to a reference ground (accessed via the green banana connector), maintaining a consistent reference potential for electrophysiology and stimulation experiments. This comprehensive grounding scheme ensures the integrity of the signal in all operating modes. All code is open-source and publicly available (see Data Availability Statement).

### Bench-top Validation

We performed two types of bench-top tests to match in-vivo stimulation parameters: measurement of signal conditioning (figure 3A) and measurement of stimulation artifacts when stimulating through an electrode and recording independently through a neurophysiology recording electrode, in a conductive saline bath (figure 3B).

**Figure 3:**
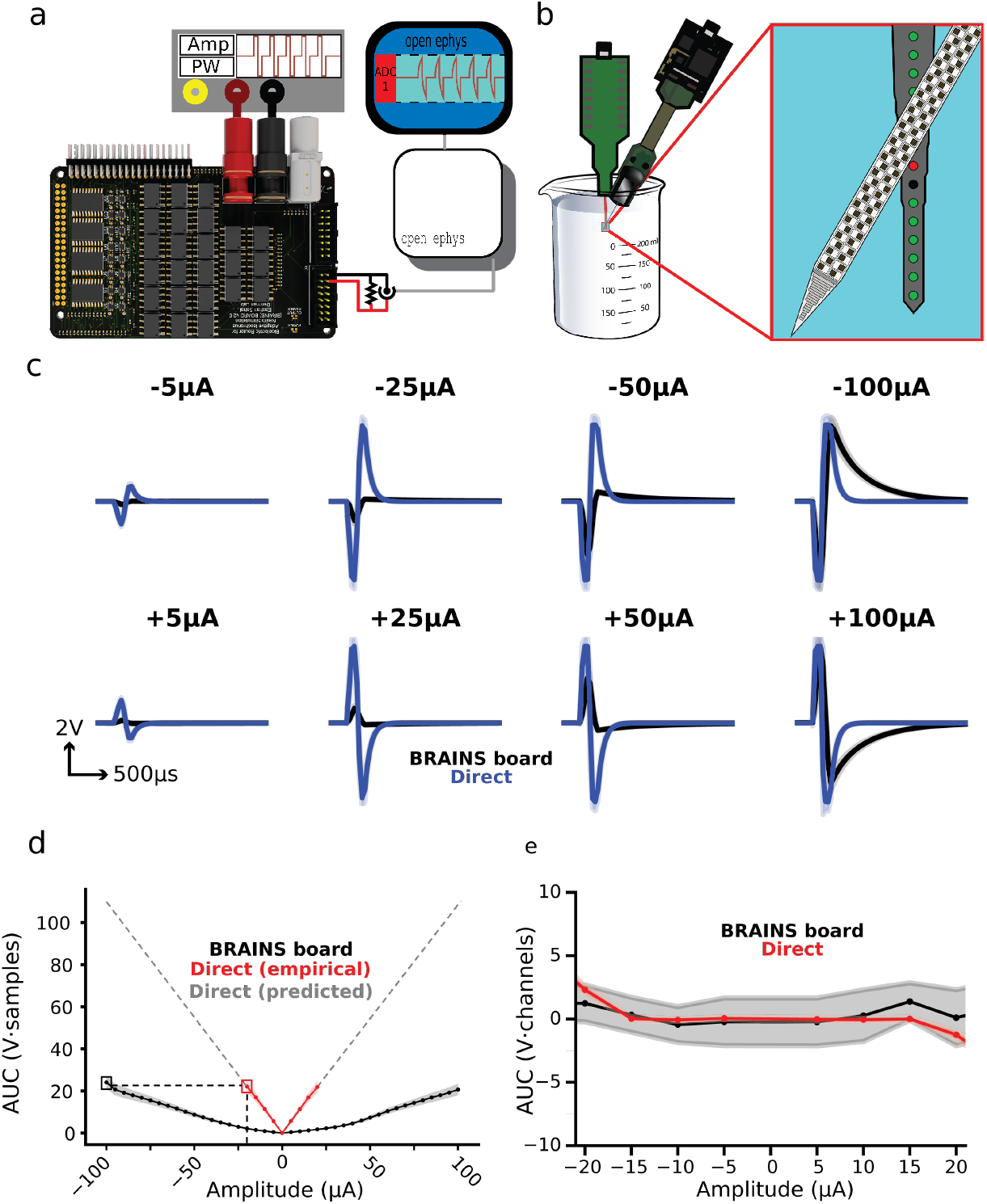
Stimulation Fidelity Bench Testing. **A**. Direct current stimulation through resistor testing setup **B**. Stimulation in saline through NeuroNexus A1×16 electrode and recording with a Neuropixel **C**. Stimulation directly through A-M Systems 4100 vs through BRAINS board with a 400kΩ resistor with varying stimuli at 100*μ*s per phase cathode and anode leading square biphasic waveforms, demonstrating the loss and shift in output waveform that is incurred due to capacitance of the solid state relay **D**. Magnitude of peak to peak area under the curve for varying amplitudes across 400kΩs for the BRAINS board and direct stimulation with a demonstration that a similar stimulus can be incurred by accounting for the capacitance by stimulating with a higher current, validated in the *in vivo* testing **E**. Peak to peak area under the curve charge balance for BRAINS board and direct stimulation parameters.

**Figure 4:**
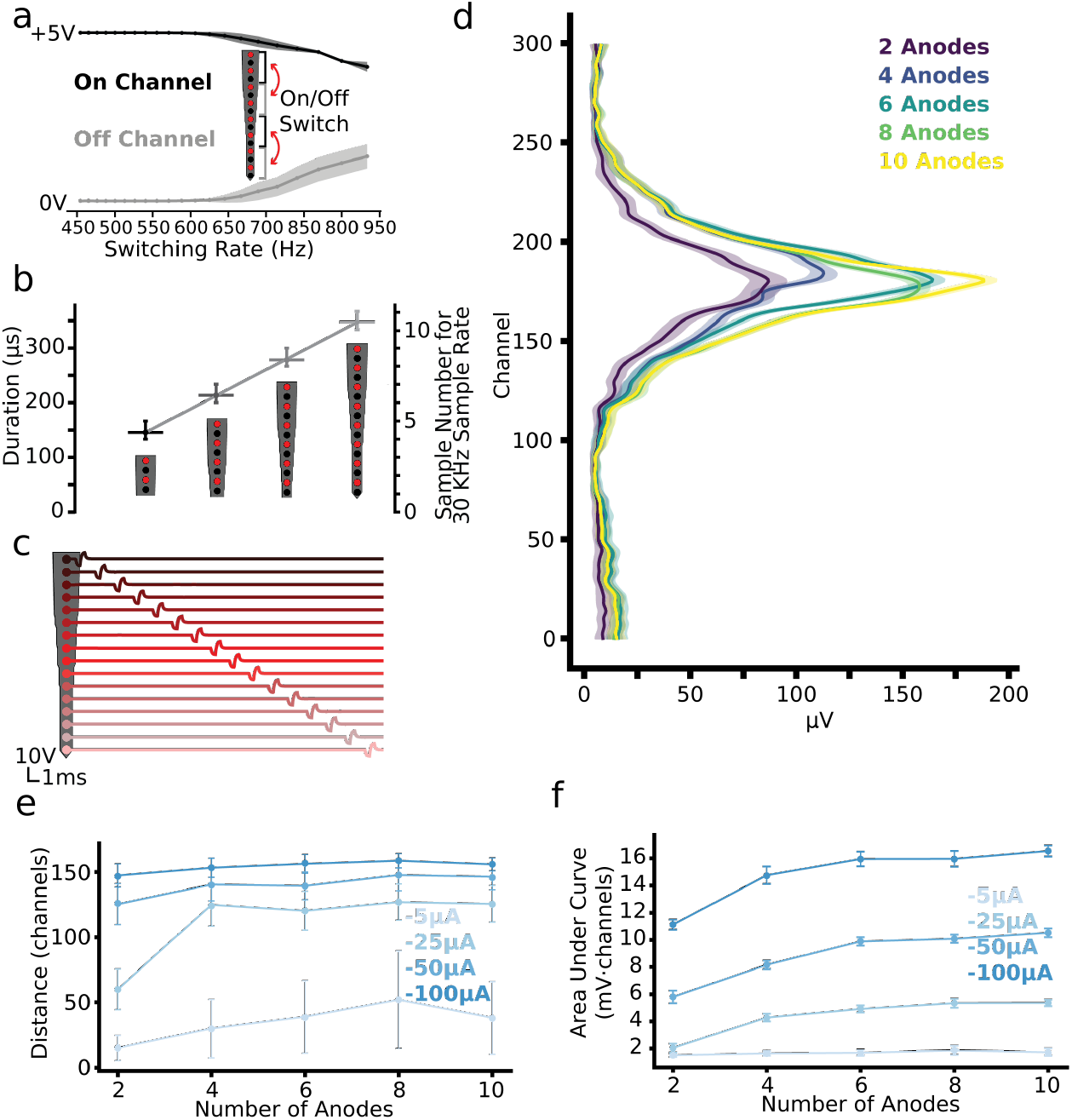
Stimulation Fidelity Bench Testing. **A**. Rapid testing (n=75 trials per frequency) of leak through off-channels and reduction of stimulus through on channels (100*μ*s per phase cathode leading biphasic stimulation at 5V amplitude through 100kΩ resistance stimulation) while switching every channels’ states from a cathode-anode pair to a ground state **B**. Output enable delay due to Software and Arduino Pro Micro delays for 1 to 4 latches enabled and switched **C**. Rapid Switching at 500Hz over 16 output monopolar channels directly connected through 100kΩ resistors with -50 *μ*A cathode leading biphasic current stimulation with 500 *μ*s waveforms with inactive channels grounded **D**. Gaussian smoothed average plot of 300 channels of a single Neuropixel recording in saline average voltage measurement over n=75 trials for -50*μ*A cathode leading biphasic stimulation through a single cathode and multiple adjacent anodes 1 ms post-stimulus delivery **E**. Average channel distance of voltage spread within a full spread threshold for Neuropixel recording of stimulation with varying amplitudes of cathode leading biphasic current stimulation through a single cathode and multiple adjacent anodes 1 ms post-stimulus delivery **F**. Average area under the curve of a 300 channel range Neuropixel recording of stimulation with varying amplitudes of cathode leading biphasic current stimulation through a single cathode and multiple adjacent anodes 1 ms post-stimulus delivery.

For signal conditioning measurements, we directly connected a high-power analog isolated stimulator (AM 4100, AM Systems) to the BRAINS board using a shielded pair of banana connectors. A third banana connector connected the signal ground of the BRAINS board to the building ground. We attached two 8-position, single-row female connectors directly to the output pins of the BRAINS board, and connected them through a resistor 10kΩ, 100kΩ, 4x 100kΩ in series, or 1MΩ), and attached a BNC connector in parallel. We attached the BNC connectors for recording into either a National Instruments PXI-6133 or an Open Ephys Acquisition Board with connections to the digital and analog I/O system. We powered and controlled the BRAINS board with an Arduino Pro Micro. Channel configurations for each experiment were set using Arduino serial commands (figure 2B). An analog isolated stimulator sent custom waveforms (constant current or constant voltage) with variable shape (anodal monophasic, cathode leading biphasic or anode leading biphasic), phase durations, amplitude, and frequency for each pulse train. Stimulator output was coupled to the BRAINS board via a resistor for each individual channel (figure 3A). A parallel set of measurements, made by recording the outputs of the analog stimulus isolator through the same resistance but without routing through the BRAINS board, served as a control for the effect of BRAINS board routing.

For measurements in saline, we stimulated through a NeuroNexus A1×16 Electrode Array in 1× Phosphate Buffered Saline (PBS) and recorded voltage with a Neuropixels 1.0 (see *In-vivo Valiation* for details on Neuropixels recording methods). We designed custom connectors using two 8-position single-row female connectors that were soldered directly to an Omnetics18 to free wire (A79045-001) connector, which directly connects to a NeuroNexus Adpt-A16-OM16 headstage. We performed impedance tests on all probes prior to experimentation using a dedicated impedance testing device (NanoZ, White Matter LLC) with an adapter to connect to the NeuroNexus array. During these tests we acquired continuous voltage series data through Neuropixels and Open Ephys GUI at 30kHz and processed it using Open Ephys python tools and custom python scripts.

### In-vivo Validation

All procedures using animals were approved by the University of Colorado Anschutz Institutional Animal Care and Use Committee (IACUC). C57BL6/J mice (n = 3) initially underwent a surgical procedure to attach an aluminum head-fixation plate to the skull. Mice were anesthetized with isoflurane (5%). The head-fixation plate was secured to the exposed skull using translucent Metabond dental cement. To seal the surgical site and facilitate later identification of lambda and bregma for sterotactic procedures, the translucent Metabond was applied to any remaining exposed skull. Following surgery, mice were given a 7-day recovery period before beginning head-fixation habituation. The habituation process involved gradually increasing the duration of head-fixation over 1-2 weeks until the mice exhibited no signs of distress during up to 2 hours of head-fixation.

Following habituation and directly before the electrophysiological recordings, mice were anesthetized and placed in a stereotaxic apparatus. Burr holes or small craniotomies were performed over the left visual cortex. Subsequently, mice were transitioned to a head-fixation platform on an in-vivo electrophysiology rig and allowed to recover from anesthesia. Neuropixels 1.0 recording electrode(s) and a multichannel stimulating electrode (Neuronexus A1×16-5mm-50-703-A16, plated with IrOx for more effective current delivery) were inserted into the brain under piezoelectrical micromanipulator control (New Scale Technologies) at a rate of 50-100μm/min to a depth of greater than 1mm. The Neuropixels recording electrodes were inserted at a 45-degree angle to the brain surface and intersected the stimulating electrode, which was inserted at a 90-degree angle (vertically through the cortical depth), approximately 100-200μm apart to prevent collisions. A stimulus isolator (AM4100, A-M Systems) was either (i) directly connected to the stimulating electrode via banana to mini-hookup clip attached to free wires from an Omnetics18 pin adapter (A79045-001, DigiKey) that mated with the A16 stimulating electrode adapter (Adpt-A16-OM16, Neuronexus) or (ii) routed through the BRAINS board via shielded banana to banana connectors. Charge-balanced bipolar and monopolar biphasic pulses with amplitudes varying from -100 to 100 μA were delivered through direct connection to stimulator and routed through the BRAINS board for direct comparison. Each parameter set was repeated 75 times with 2 seconds between pulses.

### Electrophysiological analyses

Neuropixels data was acquired using Open Ephys software. Neuropixels 1.0 implements hardware filtering on data, separating data stream into a first-order high-pass filtered (300-6000 Hz) stream (AP band) and a first-order low-pass filter (0.1-300 Hz) stream (LFP band) All data was processed and analyzed using custom Python code in Jupyter notebooks.

*RMS calculations and despiking AP band* RMS calculations were averaged from 10 1-second chunks of the AP band across each channel for both direct and BRAINS board connection setups. The voltage traces were despiked by removing values that exceeded 2.5 times the standard deviation of the mean voltage for each channel.

*LFP signal processing* 10 seconds of LFP data were extracted corresponding to each connection setup (direct and BRAINS board). These segments were baseline-corrected to remove any direct current offset by subtracting the median voltage for each channel across the selected time points. Power spectral density (PSD) analysis between 0-100 Hz was conducted using the Welch method for each channel individually. The power values were then converted to a logarithmic scale (dB). The mean gamma-band power was calculated by averaging the PSD from 30-50 Hz for each channel and smoothed for plotting with a Gaussian filter (σ= 2).

Adobe Stock Photos were utilized for the beaker in figure 3B and schematics for the Arduino setup of the BRAINS board in figure 2C were designed in Fritzing.

## Results

### Fidelity of stimulation

The fidelity of iEBS waveforms is critical to controlling the efficacy of iEBS. Any system needs to ensure high-fidelity waveforms, whether they are generated by in-built control circuitry or if the waveform passes through, as with the BRAINS board. To validate the fidelity of stimulation waveforms, we ran two types of validation tests: directly stimulating current through various resistors (figure 3A) and recorded by an analog signal acquisition system and stimulation through the stimulation electrode in saline and recording with a Neuropixel (figure 3B). There was a noticeable alteration in the signal output when routing with the BRAINS board (figure 3C), which we attribute to the output capacitance of the solid state relays (50pF, Table 2). We built an empirical comparison of the observed output magnitude to account for this capacitive drop by matching the peak to peak magnitude of current output over waveform time (figure 3D). Furthermore, the charge balance between the two devices is minimally different (p <0.001, Cohen’s d <0.2) within ranges that were recordable in direct resistor testing. (figure 3E). This is further explored with adjusted resistances of 100 kΩ and 1 MΩ resistors to demonstrate the differences between the direct current response compared to BRAINS board response in Supplemental figure 2.

### Leak and channel isolation

We designed the BRAINS board to enable multipolar stimulation through independent channel control. Our evaluation focused on characterizing the temporal dynamics and isolation properties of the stimulation channels. We demonstrated the board switches both input and output configurations across all 16 channels at 689.7 Hz with no statistically significant noise caused by latency between control output and Solid State Relay On/Off transitions (one sample t-test of the SNR of the Off Channels, p <0.1) (figure 4A). To verify the contact switching speed, we delivered electrical pulses (n=150) from an analog stimulus generator while alternating between two latch enable groups, each controlling 4 output signals. During active phases, we configured channels as either cathode or anode, defaulting to ground state when inactive, with latches 1 and 3 operating synchronously opposite to latches 2 and 4 (channel configuration details provided in figure 2A). Temporal characterization of the Arduino Pro Micro implementation revealed that the output enable signal initialization for a 4-channel configuration through a single latch required approximately 146 μs (mean=146 μs, sd=±1 sample for a 30KHz recording rate), with each additional latch or 4-channel group requiring an incremental 67 μs ±1 sample for state transitions during configuration switching (figure 4B). We did not observe significant cross-talk, with inactive (grounded) channels showing no detectable noise or cross-channel interference during rapid channel switching at 500 Hz stimulation frequencies (figure 4C). The recorded maximum and minimum voltages aligned precisely with theoretical predictions based on stimulation parameters (±5V for -50μA biphasic stimulation across 100 kΩ resistance), indicating robust channel isolation with no significant signal degradation beyond the capacitance effects previously addressed in figure 3D.

We evaluated the BRAINS board’s capacity to enable multipolar stimulation through an electrode array with a single analog stimulus isolated using a systematic characterization of observed stimulation field properties in saline. When implementing focused multipolar configurations with a single cathode and distributed anodes positioned 50 μm above and below the stimulating electrode, we observed systematic enhancement of peak voltages with increasing anode count (figure 4D). Correlation analysis revealed a strong positive relationship between anode count and maximum voltage (Spearman’s *ρ* = 0.90, p <0.05), with peak voltages increasing from 86.85 ± 3.21 μV (2 anodes) to 112.74 ± 3.06 μV (4 anodes) to 164.13 ± 3.12 μV (6 anodes) 157.88 ± 2.97 μV (8 anodes) 188.17 ± 3.07 μV (10 anodes). The spatial extent of stimulation, quantified as the full width of the voltage profile above threshold, showed a corresponding positive trend with anode count (*ρ* = 0.87, p = 0.054), expanding from 76 to 109 channels. While individual configuration comparisons did not reach statistical significance for either maximum voltage or spread width (Kruskal-Wallis test, p = 0.41 for both metrics), the monotonic relationship suggested systematic modulation of both field strength and spatial distribution through anode count manipulation.

Analysis of stimulation spread across Neuropixels recording channels (2 channels per 10 μm) revealed consistent spatial distributions across configurations, except the 2-anode, -25 μA condition (figure 4E). Statistical analysis demonstrated significant effects of both stimulation amplitude (Kruskal-Wallis H = 17.03, p <0.001) and anode count (Friedman *χ*^2^ = 13.60, p <0.01) on spread characteristics. Post-hoc analyses revealed that -100 μA stimulation produced significantly broader spreads compared to -25 μA (p <0.01) and -5 μA (p <0.001) conditions, but not -50 μA (p = 0.54). Both -50 μA and -25 μA conditions generated significantly larger spreads than -5 μA stimulation (p <0.001). While positive correlations between anode count and spread distance existed across all amplitudes (*ρ* = 0.70-0.80), these relationships did not achieve statistical significance (all p *>* 0.10). Area Under Curve (AUC) analysis across 300 channels suggested amplitude-dependent effects on total voltage distribution (figure 4F). This effect appeared most pronounced at -100 μA, where AUC values ranged from 11.1 to 16.5 mV·channels and exhibited saturation with increasing anode count. The relationship between AUC and anode count followed similar patterns across all tested amplitudes (−5 μA, -25 μA, -50 μA, and -100 μA), characterized by steep increases between 2 and 4 anodes followed by more gradual increases or plateaus. An ANOVA revealed significant amplitude effects (F = 63.14, p <0.001), with post-hoc comparisons confirming hierarchical differences between all amplitudes except -25 μA and -5 μA (p = 0.078). Linear relationships between AUC and anode count achieved significance for -50 μA and -25 μA conditions (p <0.05), while the -100 μA condition demonstrated apparent saturation, likely due to recording system limitations.

### In vivo testing

To validate the utility of the BRAINS board in experimental conditions, we inserted a linear 16-channel stimulating electrode and high-density electrophysiological recording array (Neuropixel) in mouse visual cortex (figure 5A). First, to ensure that connecting the stimulating electrode through the BRAINS board does not meaningfully increase the noise in the recording compared to direct connection, we measured the RMS of despiked high-pass voltage traces across channels (figure 5B,C). BRAINS board increased the mean RMS by 0.46 μV (direct: 10.49 μV ± 3.45, BRAINS board: 10.95 μV ± 3.14, Wilcoxon test p <0.01). This difference, while statistically significant, represents a negligible physiological difference and supports that the BRAINS board is not adding a meaningful source of electrical noise. Next, we qualitatively compared the LFP signal recorded during direct connection and connection through BRAINS board. The raw LFP signal (figure 5D), the frequency power spectra, and the gamma power across channels are remarkably similar, suggesting that connection through the BRAINS board is not altering the signal.

**Figure 5:**
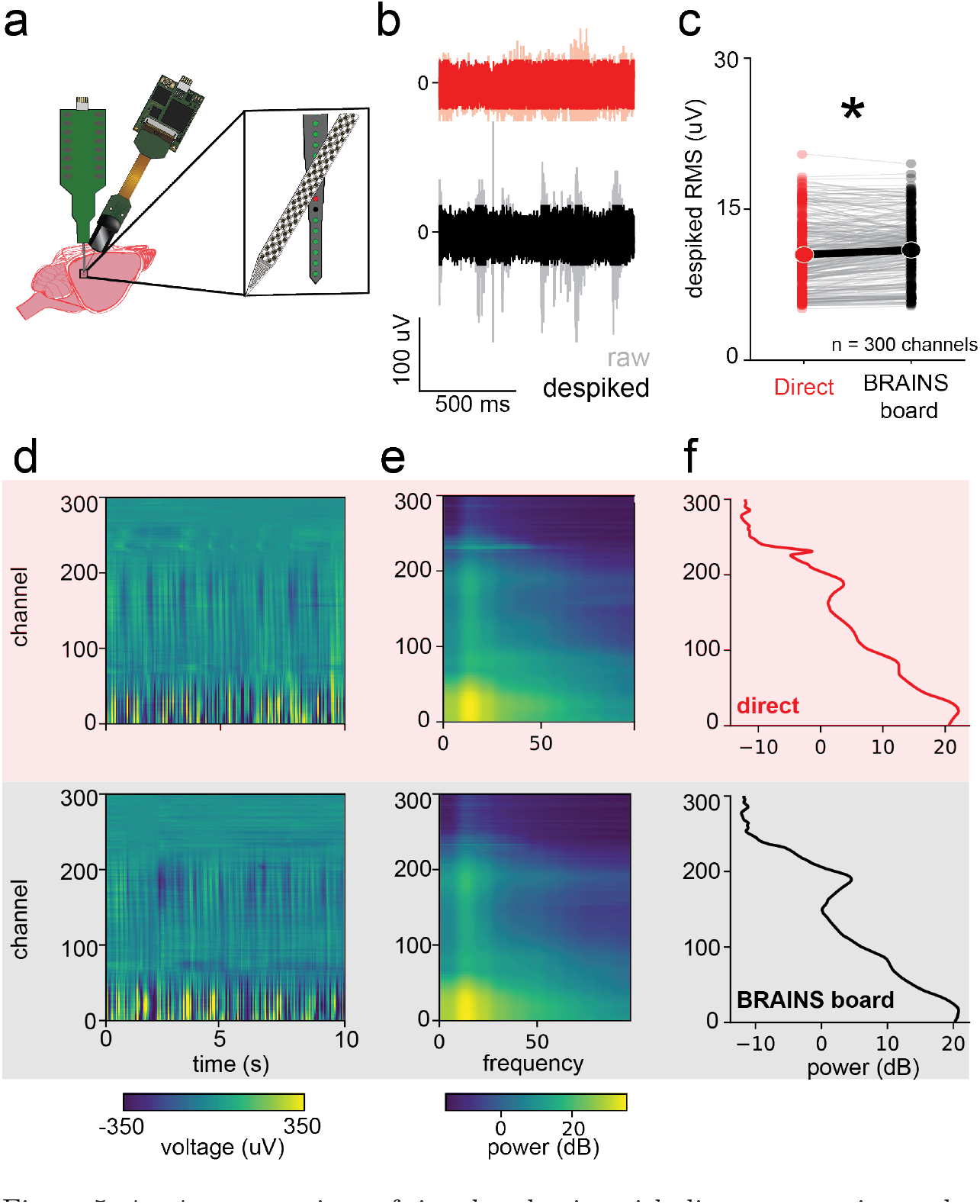
*in vivo* comparison of signal and noise with direct connection and BRAINS board. **a**. schematic for *in vivo* electrophysiology with electrical stimulation setup **b**. raw (gray and pink traces) and despiked (red and black traces) AP voltage traces from a single channel for direct connection (top, red) and BRAINS board (bottom, black). **c** paired despiked RMS with mean for direction connection and BRAINS board (n = 300 channels, Wilcoxon Test, p = 0.02). **d-f** raw LFP heatmap (d), LFP spectral power (e), and gamma power (f) for direct connection (top) and BRAINS board (bottom).

Next, we compared the stimulation efficacy through the BRAINS board compared to direct connection *in vivo*. We stimulated in mouse visual cortex while simultaneously recording the nearby artifact and evoked potential from a Neuropixel recording electrode. During prior benchtop testing, we established a near equivalent stimulation dose by using the waveform AUC measured at 400 kΩ for direct connection and BRAINS board (figure 3D). The circuit resistance changes the dose relationship (Supplemental figure 2), and we selected 400 kΩ resistance because it approximated the mean contact impedance of the stimulating electrode (407±31 kΩ), and thus an approximation of the *in vivo* circuit. Using this relationship, we identified that the measured AUC of -100 μA routed with the BRAINS board (24.054 ± 2.96) is less than 5 percent (4.8 percent) different from -25 μA direct connection (25.01 ± 2.96) in the benchtop configuration. Thus we compared -100 μA routed with the BRAINS board and -25 μA direct connection, expecting less than a 5 percent difference. The electrophysiologically measured artifact and evoked potentials at a single channel (figure 6A, top) and the corresponding voltage heatmap for all inserted recording channels (figure 6B, bottom) are visually similar for the approximated equivalent currents. To quantitatively assess similarity, we compared the AUC for the recorded extracellular voltage (including artifact and subsequent evoked potentials) for both conditions. Routing stimulation through the BRAINS board statistically increased the AUC of recorded response (direct -25 μA : 21.65 ± -2.3, BRAINS board -100 μA : 22.48 ± 1.5, unequal variances t-test: p = 0.02), but remained less than 5 percent different (4.8 percent). Thus, while we note small differences, the BRAINS board can deliver current dosages within 5 percent of those delivered through direct connection, when accounting for electrode impedances. The single channel raw voltage traces and full probe voltage heatmaps for amplitudes -100, -50, -25, -5, 5, 25, 50, and 100 for bipolar and monopolar can be viewed in Supplemental figure 3.

**Figure 6:**
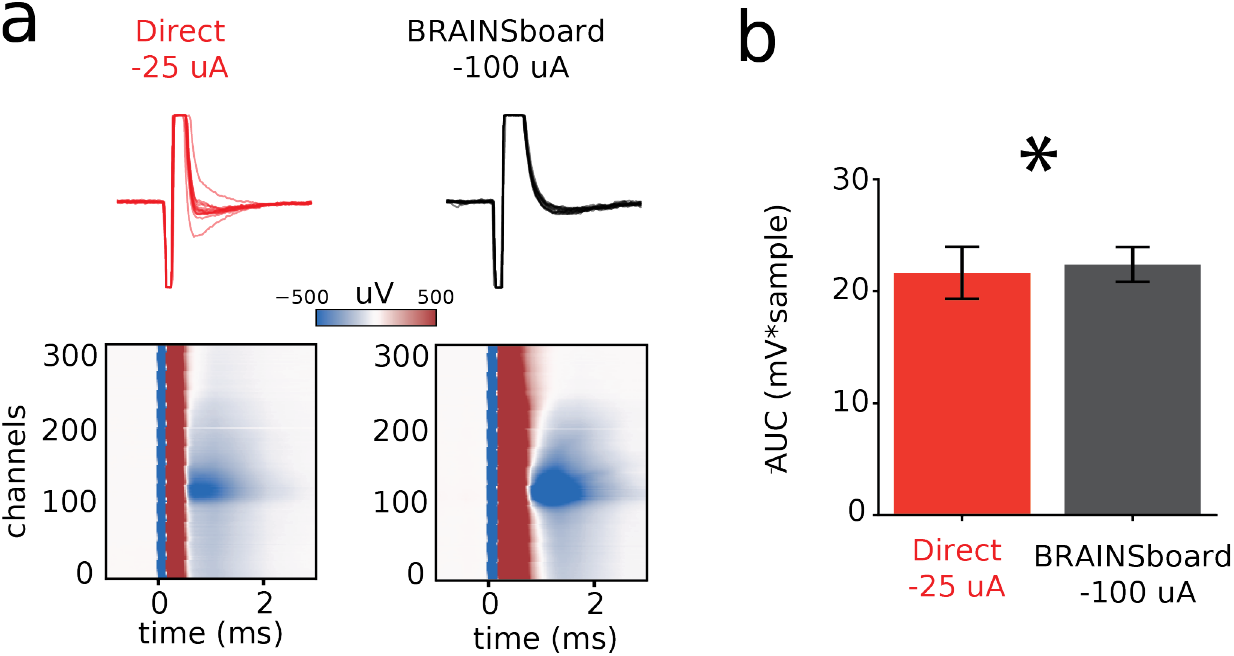
*in vivo* Comparison of BRAINS Board and Direct Connection Stimulation. **a**. single channel (channel = 150) raw voltage trace (top) and mean voltage heatmaps across all inserted channels (n = 75 trials) for matched current doses (−25 *μ*A direct and -100 *μ*A BRAINS board) measured *in vivo* in mouse visual cortex **b**. AUC for measured *in vivo* extracellular voltage during electrical pulses for -25 *μ*A direct and -100 *μ*A BRAINS board (n = 75 trials for each condition, unequal variances t-test: p = 0.02)

## Discussion

The goal of the device reported here, the BRAINS board, is to allow simultaneous multipolar and/or rapidly switching electrical stimulation through high contact count devices, particularly in research settings. We demonstrate that the BRAINS board faithfully transmits arbitrary waveforms to any available electrode (figure 2), does not introduce electrical noise to either stimulation waveforms (figure 4C) or into the nearby extracellular space, and enables both multipolar stimulation and rapid switching. The BRAINS board interfaces with any lead or passive electrode array, and we validate its use in intracortical electrical stimulation in mice. Most notably, the BRAINS board can be produced for low cost and we provide to the community specifications and parts and the open-sourced designs and software to enable manufacture of this device at or near materials cost.

Intracranial electrical brain stimulation, in both research settings (e.g., microstimulation[32, 33]) and clinical applications (e.g., deep brain stimulation[9]), is typically relies on a stimulus isolator[34]. Such isolators can be analog- or digitally-controlled, and rely on an optically-isolated battery powered circuit to generate constant voltage or constant current pulse that to a single set of anode and cathode outputs. The BRAINS board does not replace this isolator in a neuromodulation system, but instead is positioned between an isolator and the stimulus effector in tissue (i.e., the lead or electrode) to allow flexibility in the routing of the limited anode and cathode outputs of stimulus isolators. In simpler applications, the BRAINS board allows for iterative exploration of the effect of single-site or all paired bipolar configuration allowed by any electrode geometry. This functionality is particularly useful for the rapidly expanding field of high density[35, 36, 37, 38], usually silicon-based[39] neural microelectrodes. Because the BRAINS board is digitally-controlled (figure 2), when paired with a digitally-controlled stimulus isolator the spatiotemporal pattern of neuromodulation is limited only by the BRAINS board rate of switching (figure 4A-B). The BRAINS board facilitates novel application of neuromodulation through high-density electrode arrays, both extant and arising electrode technologies[40]. The BRAINS board could also be applied to empirically validate complex stimulation protocols for peripheral neuromodulation devices[**lambrecht**], where for some devices such as cochlear implants complex neuromodulation is known to have benefits[41, 42], while in others such approaches remains not yet empirically vetted (e.g., vagus nerve stimation[43]).

By enabling the routing of stimulator outputs to any connected electrode, the BRAINS board allows nearly arbitrary spatiotemporal control of neuromodulation (e.g., multipolar or temporal interference stimulation protocols) with any lead or passive electrodes. While the BRAINS board facilitates using such complex protocols with arbitrary electrodes, these protocols are not unique to the BRAINS board. One example is “current steering” through multiple contacts simultaneously[31]. The potential benefits of such protocols have been an area of active research for years. Notions of current steering for neuromodulation originated with theoretical and computational models[44, 45, 46], with the control of stimulated volume (and therefore potential limitation of off-target effects) a primary proposed benefit of current steering. However, biological validation of these proposed benefits of current-steering has been difficult. The most robust testing has come in direct clinical studies[47, 48], pre-clinical validation in animal neural tissue are limited[49]. Some pre-clincal behavioral detection studies show differences with current steering[50], but direct measurements of activated volume, and therefore insight into mechanism of action, are much more sparse[51]. The BRAINS board will allow such measurement in neurophysiology labs. to push the potential, need to try other geometries, and the BRAINS board will allow current steering with higher density and other bespoke stimulating electrodes.

In addition to current steering and directional stimulation through spatial patterning, temporal pattering is another frontier in neuromodulation[52]. Temporal patterning of single stimulation sites, where these patterns are determined *a priori*, has strong impacts on deep brain stimulation (DBS) effectiveness, with proscribed patterns able to both increase and eliminate[54] the therapeutic effectiveness. Biomimetic stimulation, where the temporal patterning is designed to mimic known statistics of the neural activity in the area being modulated, can also profoundly effect perception of neuromodulation for sensory prostheses[55]. Both *a priori* temporal patterning and biomimetic patterning could be extended to include spatiotemopral “flow” of patterns across neural circuits with multiple electrodes. Research into the effectiveness and mechanisms of these protocols is enables by the BRAINS board. Finally, “real-time” or “closed-loop”[56] control of neuromodulation based on neural or other feedback[57] to achieve an optimal modulation is a rapidly expanding form of neuromodulation. The combination of software control of rapid switching through the BRAINS board with complex neural readout available in research settings[58], can shed light on relevant biomarkers for such closed-loop neuromoduation.

The BRAINS board’s capabilities enable novel studies of electrophysiological effects of iEBS stimulation. Clinical trials have highlighted that optimal stimulation parameters may not be intuitive, and may require computational identification[59]. While clinical systems deliver directional stimulation, they cannot readily facilitate systematic investigation of neural responses, especially with the single neuron resolution across populations and neural circuits needed to optimize these therapies. Clinical research shows that LFPs features such as beta oscillations could serve as biomarkers for stimulation optimization[60]; studying how complex spatiotemporal stimulation patterns influence these population-level signals requires experimental flexibility. The BRAINS board fills this research gap by enabling precise control over stimulation parameters while allowing simultaneous electrophysiological recordings, making it possible to systematically map relationships between stimulation patterns and population dynamics. Such a systematic map, enabled by the BRAINS board, will enhance approaches to stimulation programming and could help resolve ongoing questions about how directionality, current steering, and temporal patterning influence therapeutic outcomes at the circuit level.

The BRAINS board, in its current form, enables research experiments into complex spatiotemporal neuromodulation and integration with neurophysiology tools to understand the mechanisms of neuromodulation. To continue to improve the capacities of the BRAINS board future developments the BRAINS board can be extended in future versions. Pass-through signal fidelity will be improved by upgrading the solid-state relays (e.g, to IXYS Systems, OAA160STR). This modification will reduce output capacitance from 50pF to 5pF on isolated channels and decreasing leak current from 10μA to 250nA, enhancing signal fidelity to 97% when working with high impedance electrodes. Ground loop interference remains a persistent challenge in electrophysiology experiments, particularly manifesting in the LFP band during serial command transmission. To address this, we propose integrating an embedded microprocessor directly onto the BRAINS board to establish a single ground reference point and implement comprehensive electrical isolation from computer interfaces. Alternatively, removal of Arduino dependency should notably reduce serial command-related noise in the LFP band while maintaining signal integrity across all connected components. Future development will focus on implementing modular architecture, with a base control module serving as the central processing unit and ground reference point, supplemented by attachable 16-channel shields for scalable expansion. This modular approach will facilitate integration with various neuroscience tools, including recording devices and optogenetic instruments.

In conclusion, the BRAINS board enables novel software-based and near real-time control of electrical stimulation through any stimulating electrode, facilitating the study of complex spatiotemporal pattering of intra- and transcranial neuromodulation through electrode arrays. By enabling research into such patterning with emerging research devices, the BRAINS board will advance understanding of the basic mechanisms of neuromodulation and facilitate improvements in current approaches as well as novel technologies.

## Acknowledgments

We would like to thank Grant Hughes for assistance with electrical stimulation methods used in this work and Moriah Miles for assistance with *in-vivo* testing.

## Author Contributions

Conceptualization: ES, JH, DJD

Methodology: ES, JH, DJD

Investigation: ES, JH, DJD

Visualization: ES, JH

Supervision: DJD

Writing—original draft: ES, JH, DJD

Writing—review editing: ES, JH, DJD

## Data and Materials availability

Data, software, and analysis code are available from Github, *[denmanlab/BRAINSboard](https://github.com/denmanlab/BRAINS%20board)*.

## References

[1] Benjamin Davidson et al. “Neuromodulation techniques – From non-invasive brain stimulation to deep brain stimulation”. In: Neurotherapeutics 21.3 (Apr. 2024), e00330. issn: 1878-7479. doi : 10.1016/j.neurot.2024.e00330. url: https://www.sciencedirect.com/science/article/pii/S1878747924000163 (visited on 12/11/2024).

[2] G. Darmani et al. “Non-invasive transcranial ultrasound stimulation for neuromodulation”. In: Clinical Neurophysiology 135 (Mar. 2022), pp. 51–73. issn: 1388-2457. doi: 10.1016/j.clinph.2021.12.010. url: https://www.sciencedirect.com/science/article/pii/S1388245721008920 (visited on 12/11/2024).

[3] Wynn Legon et al. “A retrospective qualitative report of symptoms and safety from transcranial focused ultrasound for neuromodulation in humans”. en. In: Scientific Reports 10.1 (Mar. 2020). Publisher: Nature Publishing Group, p. 5573. issn: 2045-2322. doi: 10.1038/s41598-020-62265-8. url: https://www.nature.com/articles/s41598-020-62265-8 (visited on 12/11/2024).

[4] John Dell’Italia et al. “Current State of Potential Mechanisms Supporting Low Intensity Focused Ultrasound for Neuromodulation”. English. In: Frontiers in Human Neuroscience 16 (Apr. 2022). Publisher: Frontiers. issn: 1662-5161. doi: 10.3389/fnhum.2022.872639. url: https://www.frontiersin.org/journals/human-neuroscience/articles/10.3389/fnhum.2022.872639/full (visited on 12/11/2024).

[5] A. J. Woods et al. “A technical guide to tDCS, and related non-invasive brain stimulation tools”. In: Clinical Neurophysiology 127.2 (Feb. 2016), pp. 1031–1048. issn: 1388-2457. doi: 10.1016/j.clinph.2015.11.012. url: https://www.sciencedirect.com/science/article/pii/S1388245715010883 (visited on 12/12/2024).

[6] Henry W. Chase et al. “Transcranial direct current stimulation: a roadmap for research, from mechanism of action to clinical implementation”. en. In: Molecular Psychiatry 25.2 (Feb. 2020), p. 397–407. issn: 1359-4184, 1476–5578. doi: 10.1038/s41380-019-0499-9. url: https://www.nature.com/articles/s41380-019-0499-9 (visited on 12/12/2024).

[7] Alessandra Del Felice et al. “Personalized transcranial alternating current stimulation (tACS) and physical therapy to treat motor and cognitive symptoms in Parkinson’s disease: A randomized cross-over trial”. In: NeuroImage: Clinical 22 (Jan. 2019), p. 101768. issn: 2213-1582. doi: 10.1016/j.nicl.2019.101768. url: https://www.sciencedirect.com/science/article/pii/S2213158219301184 (visited on 12/12/2024).

[8] A. T. Barker, R. Jalinous, and I. L. Freeston. “NON-INVASIVE MAGNETIC STIMULATION OF HUMAN MOTOR CORTEX”. In: The Lancet. Originally published as Volume 1, Issue 8437 325.8437 (May 1985), pp. 1106–1107. issn: 0140-6736. doi: 10.1016/S0140-6736(85)92413-4S0140673685924134. url: https://www.sciencedirect.com/science/article/pii/(visited on 12/12/2024).

[9] Andres M. Lozano et al. “Deep brain stimulation: current challenges and future directions”. en. In: Nature Reviews Neurology 15.3 (Mar. 2019). Publisher: Nature Publishing Group, pp. 148–160. issn: 1759-4766. doi: 10.1038/s41582-018-0128-2. url: https://www.nature.com/articles/s41582-018-0128-2 (visited on 12/12/2024).

[10] Vibhor Krishna and Alfonso Fasano. “Neuromodulation: Update on current practice and future developments”. In: Neurotherapeutics 21.3 (Apr. 2024), e00371. issn: 1878-7479. doi: 10.1016/j.neurot.2024.e00371. url: https://www.sciencedirect.com/science/article/pii/S1878747924000576 (visited on 12/11/2024).

[11] Benjamin D. Greenberg et al. “Three-Year Outcomes in Deep Brain Stimulation for Highly Resistant Obsessive–Compulsive Disorder”. en. In: Neuropsychopharmacology 31.11 (Nov. 2006). Publisher: Nature Publishing Group, pp. 2384–2393. issn: 1740-634X. doi: 10.1038/sj.npp.1301165. url: https://www.nature.com/articles/1301165 (visited on 12/06/2024).

[12] Thomas Wichmann and Mahlon R. DeLong. “Deep Brain Stimulation for Neurologic and Neuropsychiatric Disorders”. In: Neuron 52.1 (Oct. 2006), pp. 197–204. issn: 0896-6273. doi: 10.1016/j.neuron.2006.09.022. url: https://www.sciencedirect.com/science/article/pii/S089662730600729X (visited on 12/06/2024).

[13] Philippe Coubes et al. “Electrical stimulation of the globus pallidus internus in patients with primary generalized dystonia: long-term results”. eng. In: Journal of Neurosurgery 101.2 (Aug. 2004), pp. 189–194. issn: 0022-3085. doi: 10.3171/jns.2004.101.2.0189.

[14] Christianne N. Heck et al. “Two-year seizure reduction in adults with medically intractable partial onset epilepsy treated with responsive neurostimulation: Final results of the RNS System Pivotal trial”. en. In: Epilepsia 55.3 (2014). eprint: https://onlinelibrary.wiley.com/doi/pdf/10.1111/epi.12534, xpp. 432–441. issn: 1528-1167. doi: 10.1111/epi.12534. url: https://onlinelibrary.wiley.com/doi/abs/10.1111/epi.12534 (visited on 12/06/2024).

[15] Corinne Orlemann et al. “Flexible Polymer Electrodes for Stable Prosthetic Visual Perception in Mice”. en. In: Advanced Healthcare Materials 13.15 (2024). eprint: https://onlinelibrary.wiley.com/doi/pdf/10.1002/adhm.202304169, xp. 2304169. issn: 2192-2659. doi: 10.1002/adhm.202304169. url: https://onlinelibrary.wiley.com/doi/abs/10.1002/adhm.202304169 (visited on 12/06/2024).

[16] Eduardo Fernández et al. “Visual percepts evoked with an intracortical 96-channel microelectrode array inserted in human occipital cortex”. en. In: The Journal of Clinical Investigation 131.23 (Dec. 2021). Publisher: American Society for Clinical Investigation. issn: 0021-9738. doi: 10.1172/JCI151331. url: https://www.jci.org/articles/view/151331 (visited on 12/12/2024).

[17] Vernon L. Towle et al. “Toward the development of a color visual prosthesis”. en. In: Journal of Neural Engineering 18.2 (Feb. 2021). Publisher: IOP Publishing, p. 023001. issn: 1741-2552. doi: 10.1088/1741-2552/abd520. url: https://dx.doi.org/10.1088/1741-2552/abd520 (visited on 12/12/2024).

[18] A. Bolu Ajiboye et al. “Restoration of reaching and grasping movements through brain-controlled muscle stimulation in a person with tetraplegia: a proof-of-concept demonstration”. eng. In: Lancet (London, England) 389.10081 (May 2017), pp. 1821–1830. issn: 1474-547X. doi: 10.1016/S0140-6736(17)30601-3.

[19] Johannes Vorwerk et al. “A retrospective evaluation of automated optimization of deep brain stimulation parameters”. eng. In: Journal of Neural Engineering 16.6 (Nov. 2019), p. 064002. issn: 1741-2552. doi: 10.1088/1741-2552/ab35b1.

[20] Christian J. Hartmann et al. “An update on best practice of deep brain stimulation in Parkinson’s disease”. eng. In: Therapeutic Advances in Neurological Disorders 12 (2019), p. 1756286419838096. issn: 1756-2856. doi: 10.1177/1756286419838096.

[21] Günther Deuschl et al. “Deep brain stimulation: postoperative issues”. eng. In: Movement Disorders: Official Journal of the Movement Disorder Society 21 Suppl 14 (June 2006), S219–237. issn: 0885-3185. doi: 10.1002/mds.20957.

[22] J. Buhlmann et al. “Modeling of a segmented electrode for desynchronizing deep brain stimulation”. eng. In: Frontiers in Neuroengineering 4 (2011), p. 15. issn: 1662-6443. doi: 10.3389/fneng.2011.00015.

[23] Daria Nesterovich Anderson et al. “The DBS: Multiresolution, Directional Deep Brain Stimulation for Improved Targeting of Small Diameter Fibers”. eng. In: Frontiers in Neuroscience 13 (2019), p. 1152. issn: 1662-4548. doi: 10.3389/fnins.2019.01152.

[24] Simeng Zhang et al. “Comparing Current Steering Technologies for Directional Deep Brain Stimulation Using a Computational Model That Incorporates Heterogeneous Tissue Properties”. eng. In: Neuromodulation: Journal of the International Neuromodulation Society 23.4 (June 2020), pp. 469–477. issn: 1525-1403. doi: 10.1111/ner.13031.

[25] James A. Frank, Marc-Joseph Antonini, and Polina Anikeeva. “Next-generation interfaces for studying neural function”. eng. In: Nature Biotechnology 37.9 (Sept. 2019), pp. 1013–1023. issn: 1546-1696. doi: 10.1038/s41587-019-0198-8.

[26] Michael S. Beauchamp et al. “Dynamic Stimulation of Visual Cortex Produces Form Vision in Sighted and Blind Humans”. English. In: Cell 181.4 (May 2020). Publisher: Elsevier, 774–783.e5. issn: 0092-8674, 1097–4172. doi: 10.1016/j.cell.2020.04.033. url: https://www.cell.com/cell/abstract/S0092-8674(20)30496-7 (visited on 01/15/2025).

[27] Hiroshi MASUDA et al. “Surgical Strategy for Directional Deep Brain Stimulation”. In: Neurologia medico-chirurgica 62.1 (Jan. 2022), pp. 1–12. issn: 0470-8105. doi: 10.2176/nmc.ra.2021-0214. url: https://www.ncbi.nlm.nih.gov/pmc/articles/PMC8754682/ (visited on 01/08/2025).

[28] M. Fiorella Contarino et al. “Directional steering”. In: Neurology 83.13 (Sept. 2014). Publisher: Wolters Kluwer, pp. 1163–1169. doi: 10.1212/WNL.0000000000000823. url: https://www.neurology.org/doi/10.1212/WNL.0000000000000823 (visited on 01/08/2025).

[29] Thomas C. Spencer et al. “Electrical Field Shaping Techniques in a Feline Model of Retinal Degeneration”. eng. In: Annual International Conference of the IEEE Engineering in Medicine and Biology Society. IEEE Engineering in Medicine and Biology Society. Annual International Conference 2018 (July 2018), pp. 1222–1225. issn: 2694-0604. doi: 10.1109/EMBC.2018.8512473.

[30] Jessica D. Falcone and Pamela T. Bhatti. “Current steering and current focusing with a high-density intracochlear electrode array”. eng. In: Annual International Conference of the IEEE Engineering in Medicine and Biology Society. IEEE Engineering in Medicine and Biology Society. Annual International Conference 2011 (2011), pp. 1049–1052. issn: 2694-0604. doi: 10.1109/IEMBS.2011.6090244.

[31] Christopher R. Butson and Cameron C. McIntyre. “Current Steering to Control the Volume of Tissue Activated During Deep Brain Stimulation”. In: Brain stimulation 1.1 (Jan. 2008), pp. 7–15. issn: 1935-861×. doi: 10.1016/j.brs.2007.08.004. url: https://www.ncbi.nlm.nih.gov/pmc/articles/PMC2621081/ (visited on 01/08/2025).

[32] Cameron C. McIntyre and Warren M. Grill. “Selective Microstimulation of Central Nervous System Neurons”. en. In: Annals of Biomedical Engineering 28.3 (Mar. 2000), pp. 219–233. issn: 1573-9686. doi: 10.1114/1.262. url: https://doi.org/10.1114/1.262 (visited on 01/08/2025).

[33] Michael S. A. Graziano, Charlotte S. R. Taylor, and Tirin Moore. “Complex Movements Evoked by Microstimulation of Precentral Cortex”. English. In: Neuron 34.5 (May 2002). Publisher: Elsevier, pp. 841–851. issn: 0896-6273. doi: 10.1016/S0896-6273(02)00698-0. url: https://www.cell.com/neuron/abstract/S0896-6273(02)00698-0 (visited on 01/08/2025).

[34] J. Millar, T. G. Barnett, and S. J. Trout. “The neurodyne: a simple mains-powered constant-current stimulus isolator”. eng. In: Journal of Neuroscience Methods 55.1 (Nov. 1994), pp. 53–57. issn: 0165-0270. doi: 10.1016/0165-0270(94)90040-x.

[35] Jörg Scholvin et al. “Close-Packed Silicon Microelectrodes for Scalable Spatially Oversampled Neural Recording”. In: IEEE transactions on bio-medical engineering 63.1 (Jan. 2016), pp. 120–130. issn: 0018-9294. doi: 10.1109/TBME.2015.2406113. url: https://www.ncbi.nlm.nih.gov/pmc/articles/PMC4692190/ (visited on 01/08/2025).

[36] Mohammad Sadeq Saleh et al. “CMU Array: A 3D nanoprinted, fully customizable high-density microelectrode array platform”. In: Science Advances 8.40 (Oct. 2022). Publisher: American Association for the Advancement of Science, eabj4853. doi: 10.1126/sciadv.abj4853. url: https://www.science.org/doi/full/10.1126/sciadv.abj4853 (visited on 01/08/2025).

[37] Longchun Wang et al. “High-density implantable neural electrodes and chips for massive neural recordings”. en. In: Brain-X 2.2 (2024). eprint: https://onlinelibrary.wiley.com/doi/pdf/10.1002/brx2.65, e65. issn: 2835-3153. doi: 10.1002/brx2.65. url: https://onlinelibrary.wiley.com/doi/abs/10.1002/brx2.65 (visited on 01/08/2025).

[38] Zhengtuo Zhao et al. “Ultraflexible electrode arrays for months-long high-density electrophysiological mapping of thousands of neurons in rodents”. en. In: Nature Biomedical Engineering 7.4 (Apr. 2023). Publisher: Nature Publishing Group, pp. 520–532. issn: 2157-846X. doi: 10.1038/s41551-022-00941-y. url: https://www.nature.com/articles/s41551-022-00941-y (visited on 01/08/2025).

[39] Kensall D. Wise, James B. Angell, and Arnold Starr. “An Integrated-Circuit Approach to Extracellular Microelectrodes”. In: IEEE Transactions on Biomedical Engineering BME-17.3 (July 1970). Conference Name: IEEE Transactions on Biomedical Engineering, pp. 238–247. issn: 1558-2531. doi: 10.1109/TBME.1970.4502738. url: https://ieeexplore.ieee.org/document/4502738 (visited on 01/08/2025).

[40] Pingyu Wang et al. “Direct-Print 3D Electrodes for Large-Scale, High-Density, and Customizable Neural Interfaces”. en. In: Advanced Science n/a.n/a (). eprint: https://onlinelibrary.wiley.com/doi/pdf/10.1002/advs.202408602, xp. 2408602. issn: 2198-3844. doi: 10.1002/advs.202408602. url: https://onlinelibrary.wiley.com/doi/abs/10.1002/advs.202408602 (visited on 01/08/2025).

[41] Nathaniel R. Peterson, David B. Pisoni, and Richard T. Miyamoto. “Cochlear implants and spoken language processing abilities: review and assessment of the literature”. eng. In: Restorative Neurology and Neuroscience 28.2 (2010), pp. 237–250. issn: 1878-3627. doi: 10.3233/RNN-2010-0535.

[42] Jill B. Firszt et al. “Current steering creates additional pitch percepts in adult cochlear implant recipients”. eng. In: Otology & Neurotology: Official Publication of the American Otological Society, American Neurotology Society [and] European Academy of Otology and Neurotology 28.5 (Aug. 2007), pp. 629–636. issn: 1531-7129. doi: 10.1097/01.mao.0000281803.36574.bc.

[43] Chaoran Wang et al. “Vagus nerve stimulation: a physical therapy with promising potential for central nervous system disorders”. eng. In: Frontiers in Neurology 15 (2024), p. 1516242. issn: 1664-2295. doi: 10.3389/fneur.2024.1516242.

[44] Andrew P. Janson, Daria Nesterovich Anderson, and Christopher R. Butson. “Activation robustness with directional leads and multi-lead configurations in deep brain stimulation”. eng. In: Journal of Neural Engineering 17.2 (Mar. 2020), p. 026012. issn: 1741-2552. doi: 10.1088/1741-2552/ab7b1d.

[45] Ashutosh Chaturvedi, Thomas J. Foutz, and Cameron C. McIntyre. “Current steering to activate targeted neural pathways during deep brain stimulation of the subthalamic region”. eng. In: Brain Stimulation 5.3 (July 2012), pp. 369–377. issn: 1876-4754. doi: 10.1016/j.brs.2011.05.002.

[46] Jesse E. Bucksot et al. “Validation of a parameterized, open-source model of nerve stimulation”. eng. In: Journal of Neural Engineering 18.4 (Aug. 2021). issn: 1741-2552. doi: 10.1088/1741-2552/ac1983.

[47] Timo R. Ten Brinke et al. “Directional versus ring-mode deep brain stimulation for Parkinson’s disease: protocol of a multi-centre double-blind randomised crossover trial”. eng. In: BMC neurology 23.1 (Oct. 2023), p. 372. issn: 1471-2377. doi: 10.1186/s12883-023-03387-0.

[48] Akash Mishra et al. “An Institutional Experience of Directional Deep Brain Stimulation and a Review of the Literature”. In: Neuromodulation: Technology at the Neural Interface 27.3 (Apr. 2024), pp. 544–550. issn: 1094-7159. doi: 10.1016/j.neurom.2022.12.008. url: https://www.sciencedirect.com/science/article/pii/S1094715922014088 (visited on 01/09/2025).

[49] Kaviraja Udupa and Robert Chen. “The mechanisms of action of deep brain stimulation and ideas for the future development”. In: Progress in Neurobiology 133 (Oct. 2015), pp. 27–49. issn: 0301-0082. doi:10.1016/j.pneurobio.2015.08.001. url: https://www.sciencedirect.com/science/article/pii/S030100821500088X.

[50] Nicolas G. Kunigk et al. “Reducing Behavioral Detection Thresholds per Electrode via Synchronous, Spatially-Dependent Intracortical Microstimulation”. English. In: Frontiers in Neuroscience 16 (June 2022). Publisher: Frontiers. issn: 1662-453X. doi: 10.3389/fnins.2022.876142articles/10.3389/fnins.2022.876142/full. url: https://www.frontiersin.org/journals/neuroscience/ (visited on 01/09/2025).

[51] Julia P. Slopsema et al. “Orientation-selective and directional deep brain stimulation in swine assessed by functional MRI at 3T”. eng. In: NeuroImage 224 (Jan. 2021), p. 117357. issn: 1095-9572. doi: 10.1016/j.neuroimage.2020.117357.

[52] Warren M. Grill. “Temporal Pattern of Electrical Stimulation is a New Dimension of Therapeutic Innovation”. In: Current opinion in biomedical engineering 8 (Dec. 2018), pp. 1–6. issn: 2468-4511. doi: 10.1016/j.cobme.2018.08.007. url: https://www.ncbi.nlm.nih.gov/pmc/articles/PMC6426311/ (visited on 01/09/2025).

[53] Christopher W. Hess, David E. Vaillancourt, and Michael S. Okun. “The temporal pattern of stimulation may be important to the mechanism of deep brain stimulation”. In: Experimental Neurology 247 (Sept. 2013), pp. 296–302. issn: 0014-4886. doi: 10.1016/j.expneurol.2013.02.001article/pii/S0014488613000447. url: https://www.sciencedirect.com/science/ (visited on 01/09/2025).

[54] Alan D. Dorval et al. “Deep Brain Stimulation Alleviates Parkinsonian Bradykinesia by Regularizing Pallidal Activity”. In: Journal of Neurophysiology 104.2 (Aug. 2010). Publisher: American Physiological Society, pp. 911–921. issn: 0022-3077. doi: 10.1152/jn.00103.2010. url: https://journals.physiology.org/doi/full/10.1152/jn.00103.2010 (visited on 01/09/2025).

[55] Elizaveta V Okorokova, Qinpu He, and Sliman J Bensmaia. “Biomimetic encoding model for restoring touch in bionic hands through a nerve interface”. en. In: Journal of Neural Engineering 15.6 (Oct. 2018). Publisher: IOP Publishing, p. 066033. issn: 1741-2552. doi: 10.1088/1741-2552/aae398. url: https://dx.doi.org/10.1088/1741-2552/aae398 (visited on 01/09/2025).

[56] Ghazaleh Soleimani et al. “Closing the loop between brain and electrical stimulation: towards precision neuromodulation treatments”. en. In: Translational Psychiatry 13.1 (Aug. 2023). Publisher: Nature Publishing Group, pp. 1–17. issn: 2158-3188. doi: 10.1038/s41398-023-02565-5. url: https://www.nature.com/articles/s41398-023-02565-5 (visited on 01/09/2025).

[57] Roberto Guidotti et al. “When neuromodulation met control theory”. eng. In: Journal of Neural Engineering (Dec. 2024). issn: 1741-2552. doi: 10.1088/1741-2552/ad9958.

[58] James J Jun et al. “Fully integrated silicon probes for high-density recording of neural activity”. In: Nature 551.7679 (2017). Publisher: Nature Publishing Group, pp. 232–236. issn: 1476-4687.

[59] Jessica A. Karl et al. “Long-Term Clinical Experience with Directional Deep Brain Stimulation Programming: A Retrospective Review”. en. In: Neurology and Therapy 11.3 (Sept. 2022), pp. 1309–1318. issn: 2193-6536. doi: 10.1007/s40120-022-00381-5. url: https://doi.org/10.1007/s40120-022-00381-5 (visited on 01/15/2025).

[60] W. M. Michael Schüpbach et al. “Directional leads for deep brain stimulation: Opportunities and challenges”. en. In: Movement Disorders 32.10 (2017). eprint: https://onlinelibrary.wiley.com/doi/pdf/10.1002/mds.27096, xpp. 1371–1375. issn: 1531-8257. doi: 10.1002/mds.27096. url: https://onlinelibrary.wiley.com/doi/abs/10.1002/mds.27096 (visited on 01/15/2025).

