## Supplemental Figures for "A Bioelectric Router for Adaptive Isochronous Neurostimulation"

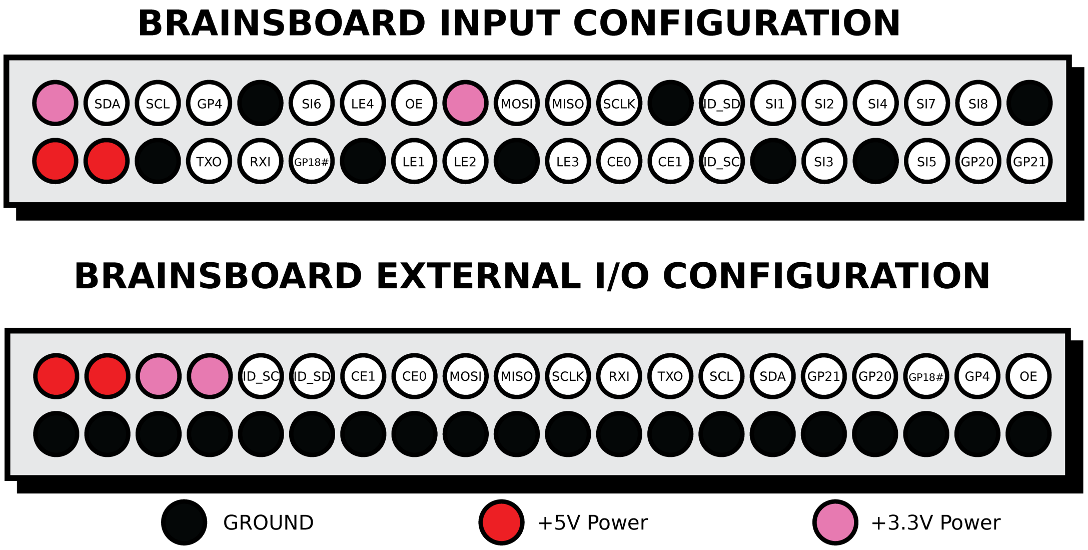


Supplemental Figure 1: BRAINS board input and output full pinout. The input configuration (top) is schematically mirrored to the pinout of a Raspberry Pi 4B, and pins that are relevant to the control are shown in figure 2D. The external I/O configuration (bottom) is on the side of the board with a 2x20 90 degree male pin header, with the entire bottom row connected to the microcontroller ground, two +5V power outputs, two +3.3V power outputs, a second connection for the OE input from the input configuration, and every unused pin by the BRAINS board itself as it relates to the input configuration.

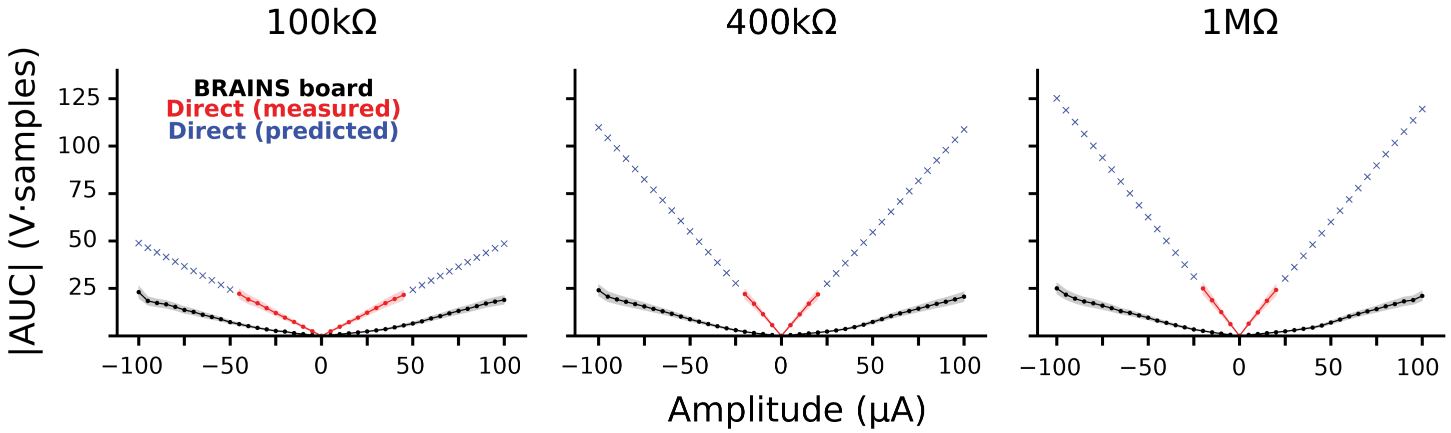


Supplemental Figure 2: A comparison between the magnitude of peak to peak area under the curve for varying amplitudes from a range of -100$\mu$A to +100$\mu$A stimulation increasing by increments of 5$\mu$A as demonstrated in figure 3D for a 400k$\Omega$ resistance. Compares the effect the capacitance has between the BRAINS board and direct stimulation for varied resistance that match high impedance electrodes, with 100 k$\Omega$ of resistance (left), 1 M$\Omega$ of resistance (right), and the original 400 k$\Omega$ of resistance (middle) (n = 75 trials per amplitude). As resistance decreases there is less of a capacitive effect and BRAINS board output will more easily match the direct current output.

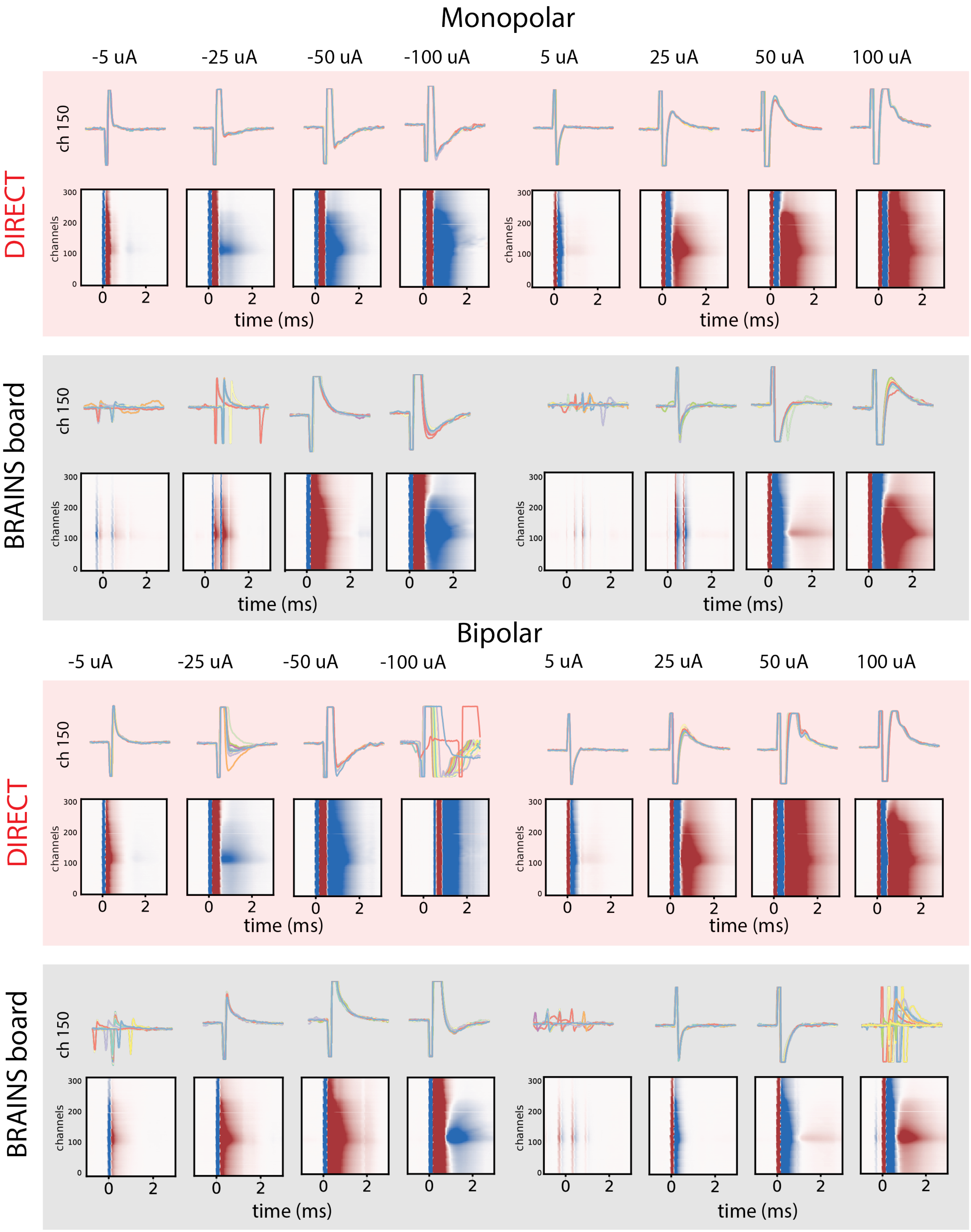
Monopolar
-5 uA
-25 uA
-50 uA
-100 uA
5 uA
25 uA
50 uA
100 uA
DIRECT
BRAINS board
ch 150
channels
time (ms)
Bipolar

Supplemental Figure 3: *in vivo* comparison of BRAINS board and direct connection stimulation for bipolar and monopolar configurations across amplitudes. Each condition contains single channel (channel = 150) voltage traces (top) and mean voltage heatmaps across all inserted channels (n = 75 trials). **first row** monopolar direct connection (red overlay) **second row** monopolar BRAINS board connection (gray overlay) **third row** bipolar direct connection (red overlay). **fourth row** bipolar BRAINS board connection (gray overlay). Trials are post-hoc aligned to stimulation onset by threshold crossing to account for trial to trial variability in stimulation onset timestamps. Some alignment failed due to response not consistently crossing threshold (see BRAINS board $\pm 5$ $\mu$A)
